# Cytokinin-mediated trichome initiation in *Nicotiana benthamiana* upon *Agrobacterium tumefaciens* infiltration

**DOI:** 10.1101/2025.11.23.690080

**Authors:** Robert Säbel, Alejandro Brand, Gerd U. Balcke, Frank Syrowatka, Claudia Horn, Sylvestre Marillonnet, Alain Tissier

## Abstract

Infiltration of *Agrobacterium tumefaciens* in leaves of *Nicotiana benthamiana* has become a standard method for transient expression in plants. Like many Solanaceae species, *N. benthamiana* develops on its leaf surface capitate glandular trichomes that produce and secrete acylsugars. We observed that *A. tumefaciens* strain GV3101 harboring the disarmed Ti plasmid pMP90 induces de novo formation of GTs and increased production of acylsugars when infiltrated in *N. benthamiana* leaves. In contrast, other strains, such as AGL1 or LBA4404, do not induce GTs. We demonstrate that the presence of the gene coding for the *trans*-zeatin synthase (*tzs*) present on pMP90 is responsible for the de novo trichome initiation and that infiltration of cytokinins such as *trans*-zeatin or benzylaminopurine is sufficient to induce GTs, establishing a direct link between cytokinins and trichome initiation. While these results provide an opportunity to better understand the development of GTs, they make us aware that Agrobacterium infiltration has developmental, biochemical and physiological consequences. To avoid or minimize potential interference of Agrobacterium on the outcome of the infiltration experiments, we recommend testing several strains when establishing a transient assay.

**Plain Language summary:** Agrobacterium infiltration of tobacco leaves is a method of choice for transient expression in plants. Agrobacterium strains that produce cytokinins induce glandular trichome initiation and acylsugar production, providing a new angle for studying trichome development and highlighting side effects of Agro-infiltration that should be considered when designing transient assays.

## Introduction

*Agrobacterium tumefaciens* is a widely used bacterial vector that mediates horizontal gene transfer and genetic transformation in plant cells. This ability relies on its tumor-inducing (Ti) plasmid that carries a transferred DNA (T-DNA) region flanked by border sequences (left and right), and a virulence (*vir*) region, encoding the type IV secretion system (T4SS) proteins, required for T-DNA processing and subsequent delivery (Lacroix and Citovsky, 2019; Weisberg et al., 2023). A wild type T-DNA from natural *Agrobacterium* species contains two classes of the so-called oncogenes. The first one encodes plant growth-promoting hormones, including auxins and cytokinins (CKs) that are responsible for the development of crown galls or hairy roots. The second one is responsible for the biosynthesis of opines, which are small molecules of amino acid-sugar conjugates that serve as carbon and nitrogen sources for *Agrobacterium* (Dessaux et al., 1998; Escobar and Dandekar, 2003). Applications in biotechnology and plant genetic transformation commonly employ disarmed *Agrobacterium* strains, in which the oncogenes inside the T-DNA region have been deleted or inactivated to prevent tumor formation. These modified Ti helper plasmids are then used in combination with a separate binary vector containing the gene(s) of interest cloned between the T-DNA borders, for use in either stable or transient transformation approaches (De Saeger et al., 2021).

Leaf infiltration of tobacco *Nicotiana benthamiana* with *A. tumefaciens* suspensions constitutes a highly efficient and time-saving method for transient gene expression assays. Commonly used disarmed laboratory strains of *Agrobacterium tumefaciens* including GV3101, AGL-1 and EHA105 are derived from the C58 strain, with the exception of LBA4404 that possesses a Ach5 chromosomal background (Hellens et al., 2000). Albeit minor, certain pathogen-related responses may alter the cell physiology of the infiltrated cells and therefore the outcome of the gene expression assays. Probably one of the best described effects on *N. benthamiana* cells after infiltration with *Agrobacterium* occurs when using GV3101 carrying the pTi plasmid pMP90. During transient assays, this strain induces the formation of stroma-filled tubules (i.e. stromules), alters plastid location around the nucleus, and promotes the accumulation of soluble sugars and starch (Erickson et al., 2014; Schattat et al., 2012). These alterations were attributed to the presence of the *trans*-zeatin synthase gene (*tzs*) in the pTi pMP90, which drives cytokinin (CK) biosynthesis upon bacterial infection (Erickson et al., 2014). Expression of the *tzs* gene results in the biosynthesis of the phytohormone *trans*-zeatin (Regier and Morris, 1982) and since the *tzs* locus is located within the *vir* region and outside the T-DNA, it is expressed but not transferred to the plant cells (Beaty et al., 1986). A direct link between GV3101(pMP90) and CKs was demonstrated when infiltration of *N. benthamiana* leaves with exogenous CKs alone mimics the changes triggered by this bacterium and its Ti plasmid (Erickson et al., 2014). Despite these side effects, GV3101(pMP90) is one of the top-performing *Agrobacterium* strains in which this CK-producing ability may have positive effects on stable transformation (Hwang et al., 2010) and can inhibit leaf senescence in infiltrated areas of tobacco (Erickson et al., 2014; Wingler and Roitsch, 2008); thereby counteracting the pathogenic effects induced by other strains over time.

In addition to the reported subcellular effects occurring shortly after GV3101(pMP90) infiltration, we also observed macroscopic changes, specifically the emergence of hairs on the leaf surface during the days following infiltration. When comparing different *A. tumefaciens* strains, including EHA105 and AGL-1, harboring the same binary vector for betalain biosynthesis, the increase in hair density is clearly visible, even to the naked eyed, and exclusively in areas infiltrated with GV3101(pMP90) after seven days (**Fig. 1a**). These hair-like outgrowths correspond to glandular trichomes (GTs) that emerge from the leaf epidermal cells and are factories of specialized metabolites. In *N. benthamiana*, capitate glandular trichomes produce acylsugars (AS) among other compounds. These sticky substances consist of a sugar core (e.g. glucose or sucrose) esterified by short- and medium chain fatty acids or acetyl groups (Ning et al., 2015; Schilmiller et al., 2012). AS are secreted from the glandular trichomes and were shown to help plants cope with biotic and abiotic stresses (Feng et al., 2022; Moghe et al., 2023; Wang et al., 2025). Trichome initiation is controlled by a combination of developmental and environmental cues through intricate molecular pathways, with phytohormones serving as key mediators of these processes (Chalvin et al., 2020; Dong et al., 2023).

**Fig. 1.**
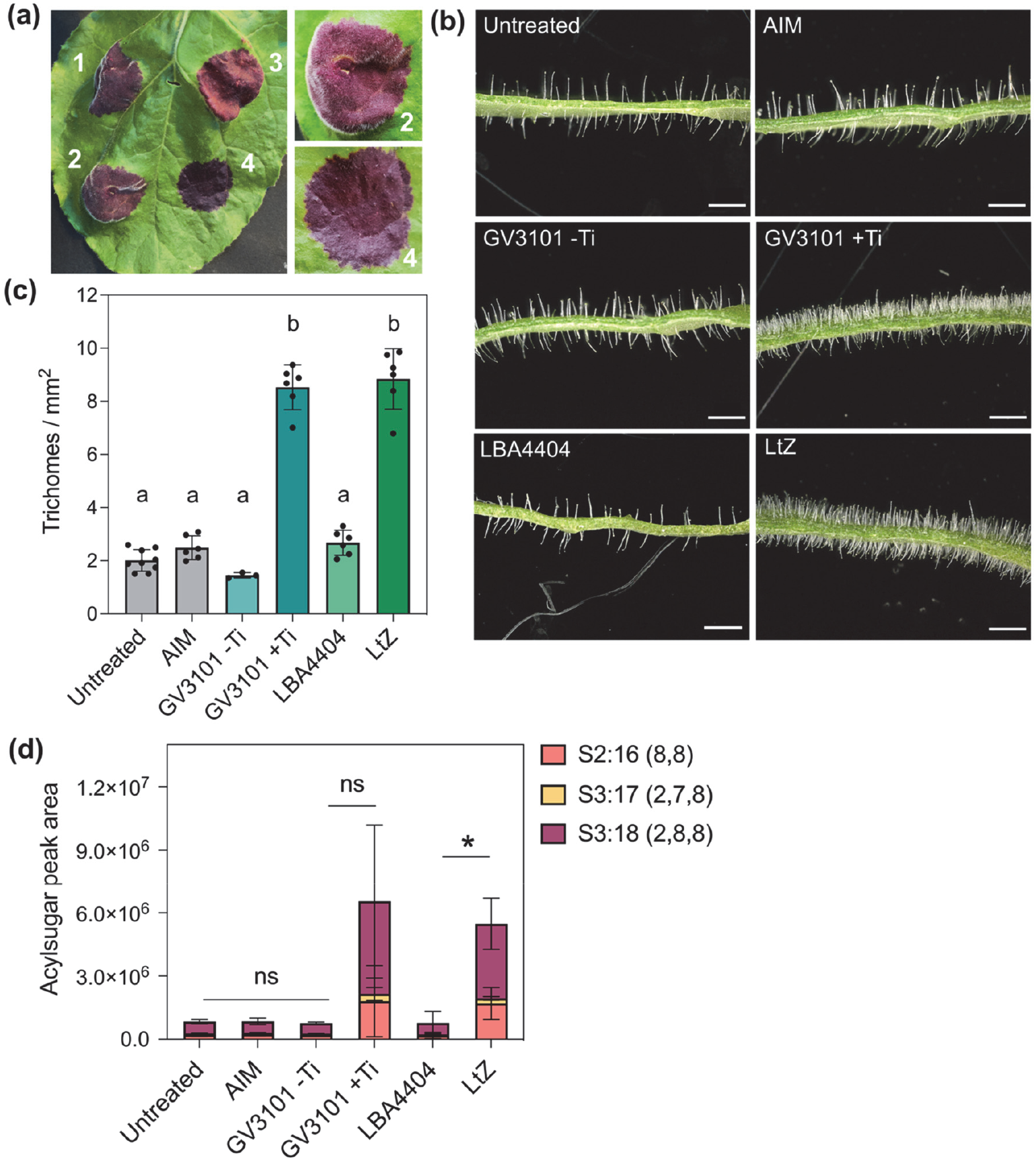
*Agrobacterium*-mediated trichome initiation in *Nicotiana benthamiana* leaves. **(a)** Leaf of an *N. benthamiana* plant infiltrated with three different *Agrobacterium* strains harboring a construct for betalain biosynthesis, four days post-infiltration (dpi). Spots 1-2: GV3101(pMP90); 3: EHA105; 4: AGL-1. The right panel shows a close-up of infiltrated spots 2 and 4, seven dpi. **(b)** Cross sections of infiltrated spots 14 dpi showing the trichomes on the adaxial and abaxial sides (see Sup. Fig. 1a, for other *Agrobacterium* strains). Abbreviations: AIM, *Agrobacterium* infiltration medium; Ti, refers to pMP90; LtZ, LBA4404 strain transformed with a vector containing the *trans-zeatin synthase* gene (*tzs*). Scale bar= 1 mm. **(c)** Trichome density on the adaxial side of infiltrated spots corresponding to the treatments depicted in (b), 10 dpi (*n* = 3-9 biological replicates). Different letters indicate significant differences following one-way ANOVA (*P* < 0.05, with Tukey’s correction) **(d)** Quantification of the main acylsugars (AS) by LC-MS/MS (*n* = 3 biological replicates). The nomenclature of AS stands as follows: S, sucrose (sugar core); followed by the number of acyl chains; the number after the colon indicates the total number of carbons in all acyl chains; and the numbers in parentheses indicate the number of carbons in each acyl chain, separated by commas. Pairwise comparisons were conducted using Student’s *t*-test, \**P* < 0.05, ns = non-significant. Error bars indicate standard deviation.

Here, we demonstrate that GV3101(pMP90) induces glandular trichomes formation in *N. benthamiana* leaves, and that this effect is mediated by transient CK production from the pTi pMP90. Infiltration with a non-CK-producing strain LBA4404 transformed with a vector carrying the *tzs* gene is capable to induce glandular trichomes. Finally, the infiltration with different CKs could mimic the induction in trichome density, accompanied by an increase in AS biosynthesis, demonstrating the direct connection of this phytohormone with trichome morphogenesis.

## Results and Discussion

To test whether trichome induction is specific to GV3101(pMP90), this strain and other common laboratory strains (LBA4404, EHA105, AGL-0 and AGL-1) were infiltrated into *N. benthamiana* leaves and trichome numbers were quantified (**Fig. 1b, Fig. S1a**). At ten days post infiltration (dpi), GV3101 carrying the pTi pMP90 (GV3101 +Ti) induced approximately threefold higher trichome density compared with untreated and control-infiltrated leaf areas (**Fig. 1b-c**). This increase in trichome density was not observed in any of the other *Agrobacterium* strains tested. After 14 dpi, some of the strains displayed yellowing of the infiltrated tissue, especially AGL-0, indicating leaf senescence. This contrasted with the long-lasting green areas infiltrated with GV3101 +Ti (**Fig. S1a**). Time-course observation of GV3101 +Ti infiltrated spots with a microscope showed a progressive increase in trichome initiation over the days after bacterial inoculation (**Fig. S1b**). Environmental scanning electron microscopy (ESEM) images revealed that trichome initiation begins as early as two dpi, marked by the bulging of pavement cells and the appearance of round glandular cells at the tip of the emerging trichomes (**Fig. S1c**). Trichome development was further assessed by quantifying acylsugars (AS). Leaf surface extracts from the infiltrated areas were analyzed by LC-MS, revealing a massive but statistically non-significant increase in AS accumulation in GV3101 +Ti-infiltrated samples, likely attributable to high biological variation (**Fig. 1d**). This result indicates that the glandular trichomes induced by *Agrobacterium* were able to produce these metabolites. To verify that trichome induction was caused by the presence of the Ti plasmid rather than the strain background, a Ti-free cured GV3101 strain (GV3101 -Ti) was generated and infiltrated following the same procedure. Both trichome number and AS levels remained comparable to the control samples (**Fig. 1b-d**). A distinct feature of the Ti plasmid of GV3101 compared with those of the other strains is the presence of the *tzs* gene in the virulence region, which is responsible for the biosynthesis of *trans*-zeatin (Beaty et al., 1986). To assess the role of the *tzs* gene and thus of CK in glandular trichome induction, we made use of a LBA4404 strain transformed with a binary vector that contains the *tzs* locus (LtZ) (Erickson et al., 2014). Leaves infiltrated with LtZ exhibited significantly more trichomes than those treated with LBA4404 and the numbers were comparable to GV3101 +Ti (**Fig. 1b-c**). These findings were also supported by the AS levels, suggesting a direct connection between trichome development and CKs (**Fig. 1d**).

The direct infiltration of synthetic CK in tobacco induces, but to a lesser extent, the same subcellular changes triggered by *tzs* from GV3101 +Ti, such as stromule formation and starch accumulation (Erickson et al., 2014). We reproduced this approach by infiltrating three different CKs: *trans*-zeatin (*t*-zeatin), kinetin and benzylaminopurine (BAP) (**Fig. 2a**). The results showed an approximately two-fold increase in trichome density, along with elevated AS levels, for *t*-zeatin and BAP (**Fig. 2b-c**). Leaves infiltrated with kinetin displayed a smaller yet significant increase in trichome number, whereas AS levels were not significantly different from the controls (**Fig. 2b-c**). In general, the infiltration with the phytohormones elicited similar responses, but with lower magnitude compared to those observed following GV3101 + Ti infiltration. This is similar to the observations of Erickson et al. (2014). One possible explanation could be the stability of the CKs inside the leaf tissue over time. For example, CK oxidase/dehydrogenases are catabolic enzymes induced upon exogenous CKs that inactivate this phytohormone as feedback mechanism to control CK homeostasis (Brugière et al., 2003; Gao et al., 2014; Schmülling et al., 2003). The fact that, in our experiments, CKs needed to be infiltrated multiple times into the same leaf area to observe the trichome phenotype highlights the importance of the continuous supply of CKs for trichome morphogenesis, which is presumably the case of the infiltrated GV3101 +Ti. Active production of *t*-zeatin by GV3101(pMP90) has been demonstrated in tissue culture experiments, where tobacco explants co-cultured with this strain were able to regenerate shoots in a phytohormone-free medium (Han et al., 2013).

**Fig. 2.**
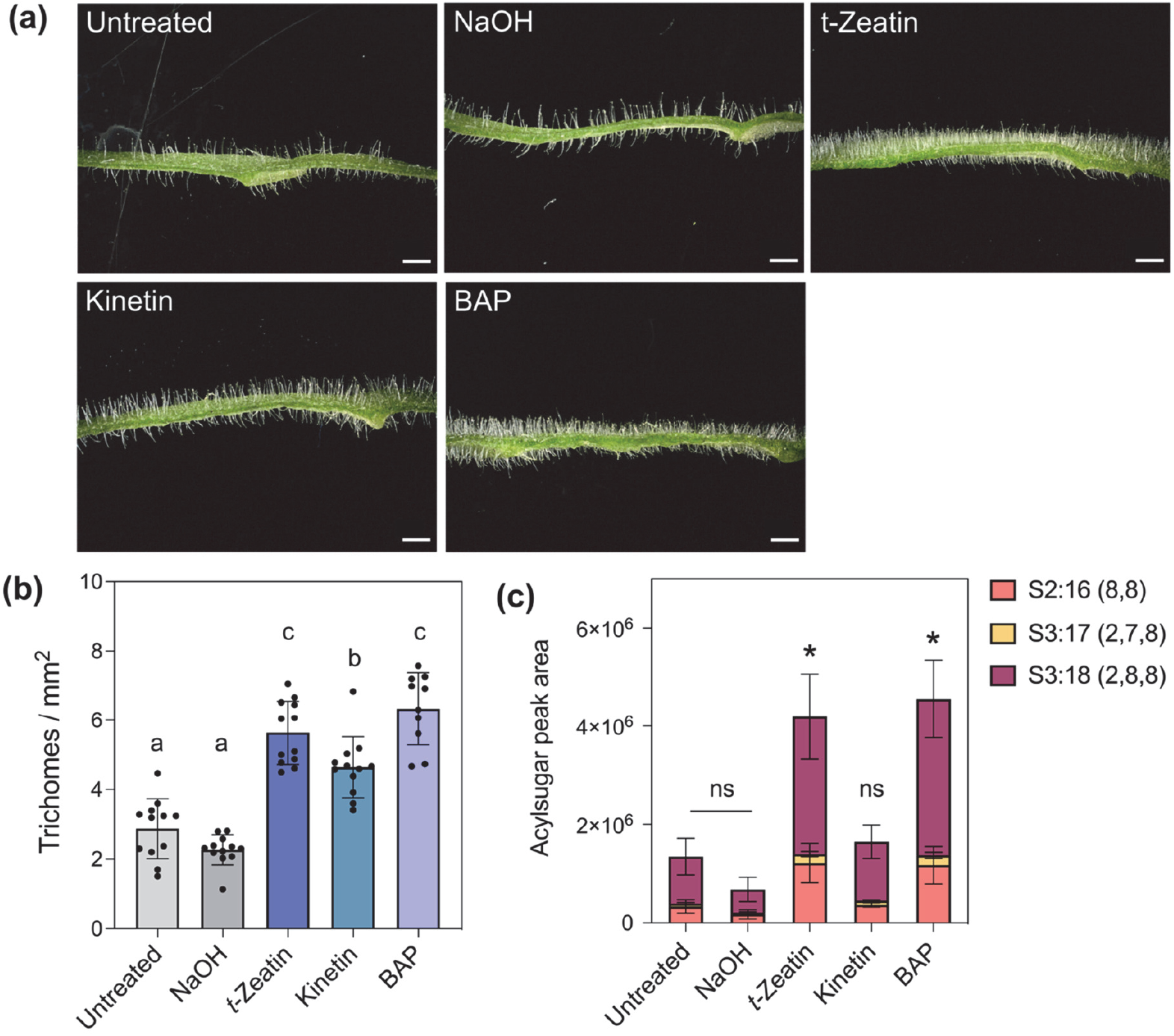
Infiltration with different cytokinins (CKs) mimics the GV3101+Ti trichome initiation in *Nicotiana benthamiana* leaves. **(a)** Effect of CKs on trichome development at 14 dpi). BAP = benzylaminopurine. **(b)** Trichome density on the adaxial side of infiltrated spots corresponding to the treatments depicted in (a) (*n* = 10-12 biological replicates). Different letters indicate significant differences following one-way ANOVA (*P* < 0.05, with Tukey’s correction). **(c)** Quantification of the main acylsugars by LC-MS/MS (*n* = 3 biological replicates). Pairwise comparisons relative to the untreated samples with Student’s *t*-test: \**P* < 0.05, ns = non-significant. Error bars indicate standard deviation. Scale bar = 1 mm.

CK is a phytohormone tightly connected with developmental processes such as cell division and leaf expansion, and cooperates with auxin to control meristem formation and maintenance (Hwang et al., 2012; Wybouw and De Rybel, 2019). Although various phytohormones contribute to trichome morphogenesis, the role of CK signaling has been comparatively less investigated than that of jasmonic acid (JA) or auxin pathways for example (Li et al., 2021). In *Arabidopsis*, which bears only non-glandular trichomes, expression of the CK biosynthesis gene *isopentenyl transferase* (*IPT*) from *Agrobacterium* in the carpel leads to ectopic trichome development (Greenboim-Wainberg et al., 2005). In tomato, which produces both glandular and non-glandular trichomes, exogenous BAP specifically induced glandular trichomes, including uni- and multicellular head types, whereas JA exhibited a broader effect on all trichome types (Maes and Goossens, 2010). Similar results were shown for *Artemisia annua*, where BAP-treated plants exhibited more glandular and filamentous trichomes, although with reduced gland size and producing lower levels of the antimalarial compounds (Maes et al., 2011). These reports illustrate the distinct regulatory mechanisms that CKs and other phytohormones exert over different trichome types and their metabolism across plant species.

Most transcription factors (TFs) implicated in CK-mediated trichome development have been characterized in *Arabidopsis* (reviewed in Li et al. (2021)). For example *GLABROUS INFLORESCENCE STEMS 2* (*GIS2*) and *ZINC FINGER PROTEIN 8* (*ZFP8*), are two C2H2 zinc-finger TFs that positively regulate trichome initiation on stems and flowers of *Arabidopsis*, through crosstalk between CKs and gibberellins (GA) (Gan et al., 2007). To the best of our knowledge, molecular factors regulating trichome development associated with CKs have not been characterized in tobacco and other species that have glandular trichomes. *NbGIS* TF, homolog of *AtGIS*, promotes trichome initiation through GA signaling by activating *NbMYB123-like* TF (Liu et al., 2018). Although *AtGIS* operates independently of CK signaling, it is still unknown whether *NbGIS* or homologs of the *Arabidopsis* C2H2 zinc-finger TFs play conserved functions in CK responses and glandular trichome initiation in tobacco.

The CK-mediated trichome induction observed upon infection with *Agrobacterium* GV3101(pMP90) reveals additional effects of using this strain, besides the previously described sub-cellular changes (Erickson et al., 2014; Schattat et al., 2012). CKs derived from bacteria are likely to trigger transcriptional changes associated with trichome development, and these effects should be considered when interpreting transient expression assays. Trichome initiation typically begins in young, developing leaf primordia at early stages of epidermal cell differentiation (Chopra et al., 2019; Larkin et al., 1996). The massive trichome induction recorded upon CK infiltration hints at a role for CKs in cellular reprogramming and in maintaining the competence of epidermal cells for trichome morphogenesis. This raises several questions. For example, whether the increase in trichome density is accompanied by *de novo* cell division or instead relies on the acquired competence of existing cells to develop these hair structures. Overall, this phenomenon, perhaps unnoticed for a long time since the widespread adoption of GV3101(pMP90) in many laboratories, provides a novel system for investigating the molecular factors that control cell fate and the role of CKs in trichome developmental and secretory processes. In addition, as infiltration of *N. benthamiana* by *Agrobacterium* is widely used in the plant community for a variety of assays, ranging from the expression of biosynthesis pathways to that of disease resistance genes, we recommend checking whether CKs, the presence of glandular trichomes or of AS could influence the outcome of the assay. This can be simply done by comparing infiltration with GV3101(pMP90) and a strain which does not produce CKS, such as LBA4404.

## Acknowledgements

The authors thank Dr. Jessica Erickson for kindly providing the *Agrobacterium* LtZ strain. This work was funded by in part by the project *KETCHUP* of the Leibniz Association (Project number K287/2019) and core funds of the Leibniz Institute of Plant Biochemistry.

## Author contributions

RS performed most of the experiments and analyzed the data, AB wrote the manuscript draft performed some experiments, GUB measured the AS, FS performed the ESEM, CH generated the cured GV3101 Agrobacterium strain and conducted the infiltration of the vectors for betalain production, SM first observed the trichome phenotype and designed experiments, AT acquired funding, supervised the project and corrected the manuscript. All authors read and approved the manuscript.

## Supplementary Information

**Supplemental Figure S1.**
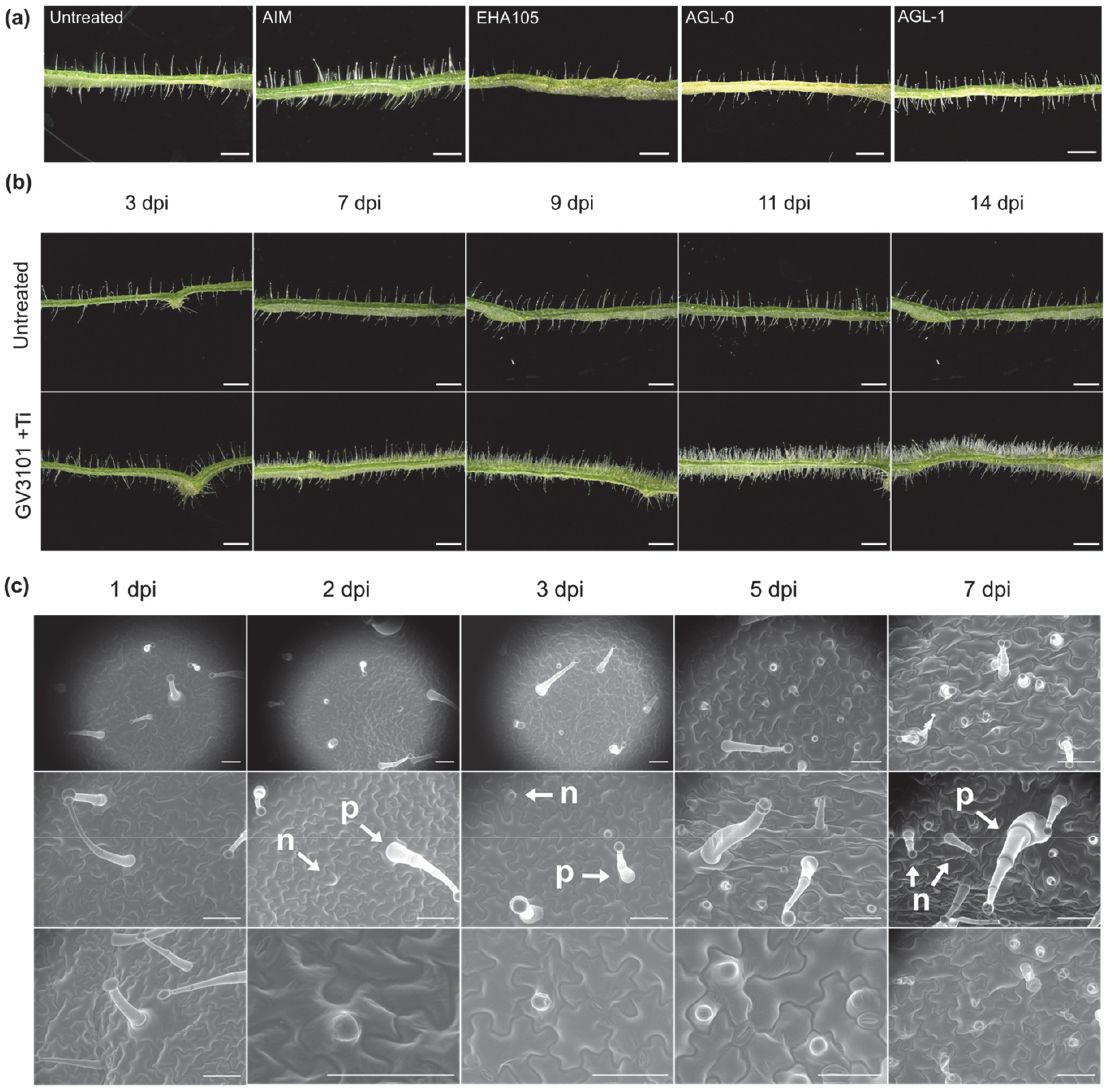
**(a)**Trichome development on the adaxial and abaxial sides of leaves of *Nicotiana benthamiana* infiltrated by different *Agrobacterium tumefaciens* strains 14 days post-infiltration (dpi). AIM: Agrobacterium infiltration medium. **(b)** Trichome density over time after infiltration with GV3101(pMP90). Scale bar = 1 mm. Ti, refers to pMP90 **(c)** Environmental scanning electron microscopy images of trichome initiation induced after infiltration by GV3101(pMP90). p: preexisting trichomes; n: newly emerging trichomes. Scale bar = 100 µm.

## Materials and methods

### Agrobacterium strains

*Agrobacterium* strains GV3101(pMP90), LBA4404, EHA105, AGL-0, AGL-1 were obtained from commercial providers. The pMP90 Ti plasmid was eliminated from GV3101 following the protocol of Hynes et al. (1985). Briefly, the GV3101 bacteria harboring the Ti plasmid pMP90 were grown on LB agar medium containing 50 mg ml^-1^ Rifampicin for three days at 40°C, followed by two days at 28°C. Single colonies were streaked onto both Rifampicin-Gentamicin and Rifampicin-only plates. The LBA4404 strain, transformed with a binary vector containing the *trans-zeatin synthase* gene (*tzs*) was kindly provided by Dr. Jessica Erickson (IPB, Halle) (Erickson et al., 2014).

### Infiltration of N. benthamiana leaves

Different *Agrobacterium* strains were grown on LB agar medium containing the respective antibiotics for two days and resuspended in *Agrobacterium* infiltration medium (AIM: 10 mM MES, 10 mM MgCl_2_, 10 µM acetosyringone). Cultures were adjusted to OD_600_ of 0.2 and incubated for two hours at room temperature. Cultures were then infiltrated into leaves of five-week-old *Nicotiana benthamiana* plants. For cytokinin infiltration, each phytohormone (*trans*-Zeatin, kinetin, and 6-benzylaminopurine (BAP), from Duchefa Biochemie B.V. Haarlem, The Netherlands) was first dissolved in 0.5 mL of 1 N NaOH and subsequently in water for 100 µg ml^->1^ >working solutions. Plants were infiltrated three times, every other day. The vector pAGM50583 for betalain biosynthesis (Grützner et al., 2021) was used to help visualize the trichomes, which appear white on a red background.

### Trichome counting and imaging

Leaf discs were excised from infiltrated spots using a cork borer (diameter 1 cm). Trichomes were manually counted on 1-cm diameter leaf discs with the VHX-6000 microscope in combination with a VH-Z20 T zoom lens (Keyence, Osaka, Japan). Images shown in Figures 1-2 were obtained by cutting 1.5 mm wide leaf strips and rotating them 90°. Environmental scanning electron microscopy images were recorded using an ESEM XL-30 FEG (FEI/Philips, Eindhoven, The Netherlands) at specified time points after agroinfiltration.

### Acylsugar profiling

Leaf surface extracts were obtained by dipping one 1-cm-diameter leaf disc into 0.5 mL of 80% methanol solution and inverting the tubes for 2 min. Representative acylsugars (AS) produced by *Nicotiana benthamiana* glandular trichomes were quantified by LC-MS/MS. Metabolites were separated with an Acquity UPLC (Waters Inc.) with a Nucleoshell RP18 column (Macherey & Nagel, 150 mm × 2 mm × 2.7 μm). Solvents and gradient were adjusted according to Balcke *et al*. (2024). Mass spectrometric analysis was performed by MS1 full scan from 65-1500 Daltons and 100 ms accumulation time (ZenoToF 7600, Sciex GmbH, Darmstadt, Germany) operating in negative mode and controlled by Sciex OS software (Sciex). The identity of specific AS was confirmed by MS/MS spectral matching to an inhouse library. MS1 AS peak areas were determined with Multiquant 3 software (Sciex GmbH, Darmstadt, Germany).

